# Fast alignment of mass spectra in large proteomics datasets, capturing dissimilarities arising from multiple complex modifications of peptides

**DOI:** 10.1101/2023.03.09.531667

**Authors:** Grégoire Prunier, Mehdi Cherkaoui, Albane Lysiak, Olivier Langella, Mélisande Blein-Nicolas, Virginie Lollier, Emile Benoist, Géraldine Jean, Guillaume Fertin, Hélène Rogniaux, Dominique Tessier

## Abstract

**Background:** In proteomics, the interpretation of mass spectra representing peptides carrying multiple complex modifications is still challenging, currently limited by the number of potential modifications considered in a single analysis and the need to know them in advance. Further developments must be done in the field to help the scientific community to discover new post-translational modifications that play an essential role in disease and to understand how chemical modifications carried by food proteins could impact our health.

**Results:** To make progress on this issue, we implemented SpecGlobX (SpecGlob eXTended to eXperimental spectra), a standalone Java application that quickly determines the best spectral alignments of a (possibly very large) list of Peptide-to-Spectrum Matches (PSMs) provided by any open modification search method, or generated by the user. As input, SpecGlobX reads a file containing spectra in MGF or mzML format and a semicolon-delimited spreadsheet describing the PSMs. As output, SpecGlobX returns the best alignment for each PSM, splitting the mass difference between the spectrum and the peptide into one or more shifts while considering the possibility of non-aligned masses (a phenomenon resulting from many situations including neutral losses).

SpecGlobX is fast, able to align one million PSMs in about 1.5 minutes on a standard desktop. Firstly, we remind the foundations of the algorithm and detail how we adapted SpecGlob (the method we previously developed following the same aim, but limited to the interpretation of perfect simulated spectra) to the interpretation of imperfect experimental spectra. Then, we highlight the interest of SpecGlobX as a complementary tool downstream to three open modification search methods on a large simulated spectra dataset. Finally, we show on a smaller dataset that SpecGlobX performs equally well on experimental and simulated spectra.

**Conclusions:** SpecGlobX is helpful as a decision support tool, providing keys to interpret peptides carrying complex modifications still poorly considered by current open modification search software. Better alignment of PSMs enhances confidence in the identification of spectra provided by open modification search methods and should improve the interpretation rate of spectra.

## Background

Interpreting fragmentation mass spectra and their assignment to a peptide sequence is an important and challenging issue in proteomics, especially when peptides carry one or several modifications. These modifications could explain the low rate of interpreted spectra after a standard bottom-up mass spectrometry analysis [1, 2]. While the modifications on proteins may be the consequence of co- or post-translational events in the cell, they may also be due to sample preparation (use of chemicals during the preparation, e.g., surfactants), to the presence of allelic variants not described in the protein databases, or may be induced by protein processing during the preparation of foods (such as Maillard’s reaction). Although reference databases of modifications exist and can draw up a list of more than a thousand modifications such as in Unimod [3], the variety of putative modifications is so large that existing databases cannot be exhaustive. Besides, it is also difficult, if not impossible, to predict and account for all possible modifications occurring in a sample.

In recent years, several open modification search (OMS) methods have been proposed to enhance the identification of mass spectra corresponding to modified peptides. These identifications are achieved through the comparison of experimental spectra to ‘reference spectra’, either simulated *in silico* from peptide candidates - referred to as theoretical spectra - or collected in a spectral database [4]. Whereas conventional methods (also called restricted or closed-search methods) try to identify experimental spectra in comparison to reference spectra in a narrow search mass window, OMS methods evaluate all or at least a large part of the reference spectra without any (or only a limited restriction) on their masses. Several advantages distinguish OMS methods from their conventional methods counterparts: (1) they enable the interpretation of spectra corresponding to peptides with unanticipated modifications; (2) the large number of modifications they can underpin does not alter identification results confidence as it occurs when an extremely large search space is generated by the introduction of many variable modifications on simulated spectra in the reference spectra database [5, 6].

The result of an OMS method is a list of PSMs (*St*, *Se*, ΔM) where *St* denotes the theoretical fragmentation spectrum of a candidate peptide, *Se* denotes an experimental spectrum, and ΔM is the mass difference between *St* and *Se*. This mass difference ΔM, if above the mass accuracy of the instrument, should be explained by one or several modifications carried by the peptide in the experience. Therefore, one or more mass shift(s) (resulting from the modification(s)) should be applied to *St* to align it with *Se*. When *Se* displays the fragmentation pattern of a peptide carrying only one modification, identification and localization (more or less precisely according to the software) of this modification are already resolved by several methods [7, 8, 9, 10, 11], mainly by testing successively the location of the modification on each amino acid. The detection of pairs of modified and unmodified peptides that may coexist in the same sample can also boost the sensitivity of the protein modification mapping [12]. When *Se* corresponds to a peptide carrying more than one modification, the interpretation is much more complicated because ΔM has to be split into several mass shifts. Several software [13, 14, 15, 16, 17, 18] attempt to explain ΔM by a combination of masses related to modifications stored in a predetermined list, or deduced from the most frequent ones observed in the sample. However, to our knowledge, there is currently no open-source software adapted to routine laboratory use allowing the ΔM interpretation without any *a priori* on the modifications, including labile ones. To fill this gap, we implemented SpecGlobX, which aligns very quickly pairs of spectra (*St*, *Se*).

SpecGlobX (SpecGlob eXtended version adapted to eXperimental spectra) is a standalone software based on a dynamic programming algorithm. It derives from SpecGlob [19], which we first developed using perfect simulated spectra. This initial step was essential to evaluate the SpecGlob algorithm’s ability to consider complex and variable modifications and to optimize its execution time. To reach a fast execution speed, we introduced - among other things - a simplification in the alignment of spectra compared to previous methods also based on dynamic programming [20, 21, 22]: each peptide is modeled by a series of peaks corresponding to only the b-ions generated by an ideal fragmentation. However, on their side, experimental spectra are not ideal; they lack some fragmentation peaks and are noisy. In the present document, we present the main adaptations developed in SpecGlobX towards the applicability of SpecGlob on experimental spectra, considering the imperfections of these spectra. Then, we highlight the benefits that SpecGlobX delivers downstream to three OMS methods: SpecOMS [23], MODPlus [14], and MSFragger [9] on a simulated dataset that mimics experimental spectra. Indeed, even if the complexity of an experimental spectrum is difficult to reproduce, simulated spectra are helpful to understand the behavior of an algorithm when it is difficult - if not impossible - to be sure of the large-scale interpretation of PSM lists. Finally, we challenged SpecGlobX on a set of experimental spectra already interpreted by different OMS software, namely the HEK293 dataset [24]. Overall, we demonstrate that SpecGlobX highlights modifications that other methods missed while maintaining a good execution time.

## SpecGlobX implementation

The algorithm of SpecGlobX follows three processing steps described in detail below: first, the completion of experimental spectra compensates for missing peaks; second, the alignment between completed experimental spectra and theoretical spectra including possible mass shifts; third, the optimization of the location of the suggested mass shifts (post-processing step).

### Completion process of the experimental spectra

Usually, a peptide *p* with an amino acid sequence *a_1_a_2_…a_i_…a_n_* is represented by a spectrum containing peaks that correspond to the b-ions {b_1_,…, b_i_,…, b_n_} and the y-ions {y_1_,…, y_i_,…, y_n_} generated by a perfect fragmentation. Then, a mass modification applied to amino acid a_i_ results in a mass shift of the peaks b_i_ to b_n_ and y_1_ to y_n-i+1_, all the remaining peaks being unchanged. The shift of only a part of the peaks creates a complex situation to manage when aligning spectra with a dynamic programming approach. To overcome this difficulty, in SpecGlobX, a peptide *p* is only represented by its b-ion peaks, so spectra alignment only requires aligning pairs of b-ion peaks from *St* (representing amino acids) with pairs of b-ion peaks from *Se*. However, distinguishing b- from y-ions in an experimental spectrum is not easy *a priori* (i.e., before interpretation). Moreover, it is well known an experimental spectrum is likely to have several missing b-ion peaks. However, SpecGlobX needs as many b-ion peaks as possible to adjust its alignment. So, as we did in [25], SpecGlobX transforms *Se* into a new spectrum called *Se_c_* (for completed *Se*) by adding new peaks according to the following rule: each original peak in *Se* is hypothesized at first as a b-ion peak (assumption 1), so it can be directly compared to the b-ion peaks of the theoretical model (*St*); then each original peak in *Se* is hypothesized as corresponding to a y-ion (assumption 2); in that latter situation, SpecGlobX adds a new peak in the spectrum *Se_c_*, called a “complementary peak”, whose mass is M + m - mass(peak), where M is the total mass of the experimental spectrum *Se* and *m* the mass of the proton. We underline that a complementary peak in *Se_c_* can replace a missing b-ion peak provided the y-ion peak is observed in *Se*. When a peptide fragmentation produced both b- and y-ions simultaneously in *Se*, except in particular situations we will focus on later in this manuscript, the “completion process” does not add any peak. Conversely, if the fragmentation process has generated only one of the two expected ions, an additional peak is added in *Se*_c_ compared to *Se*. Given the accuracy of fragment masses in current mass spectrometers (of the order of 0.005 Da), we anticipate that the noise added by complementary peaks should not significantly interfere with alignments. The generation of a series of complementary peaks is illustrated in Figure 1c.

**Figure 1.**
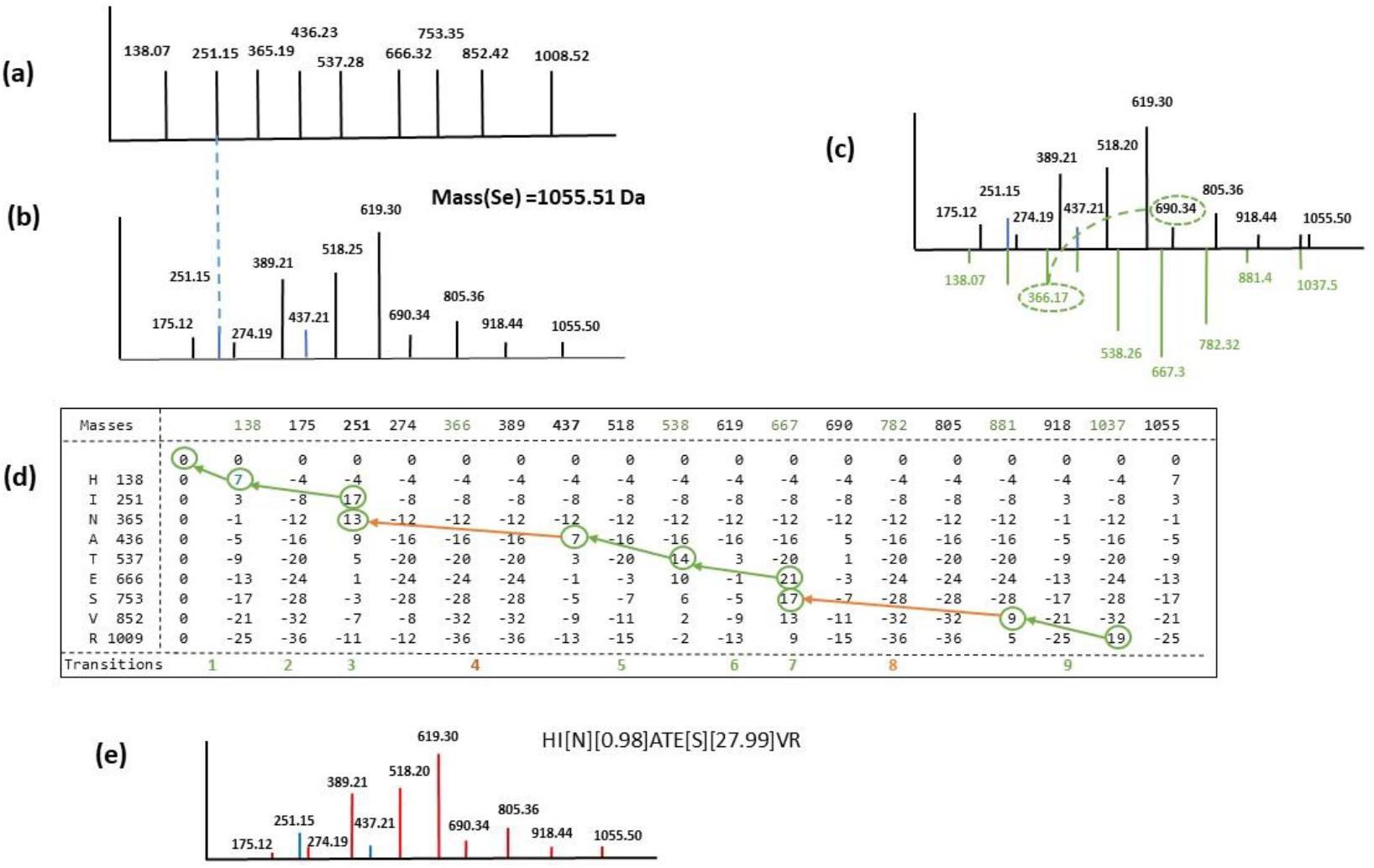
Alignment of peptide HINATESVR on a simulated spectrum carrying two modifications. (a) The theoretical spectrum, *St*, of peptide HINATESVR modeled only by its b-ion peaks. (b) The simulated spectrum, *Se*, with dummy intensities (intensities have no effect during the alignment with SpecGlobX). *Se* contains two modifications compared to *St* and only two b-ion peaks (represented in blue) are present in the spectrum. (c) Complementary peaks – below the x-axis - are added to *Se*, to generate the spectrum *Se_c_*; For example, one ‘original peak’ and its complementary peak are connected by a dotted line for better readability. (d) The score matrix was computed by SpecGlobX to align both spectra by dynamic programming. Masses have been rounded for simplification of the graphical representation; masses corresponding to complementary peaks are in green; transitions are numbered on the last row. The green (when SpecGlobX found an alignment) or orange arrows -when SpecGlobX found a realignment (with a mass shift)-delineates the path followed by the traceback step interpreting *Se*. (d) The interpretation of *Se* as HI[N][0.98]ATE[S][27.99]VR in terms of peptide sequence and mass shifts suggest a deamidation on “N” (mass increment of 0.98 Da) and a formylation on “S” (mass increment of 27.99 Da); non-aligned amino acids and mass shift values are in brackets. Original b-ion peaks are in blue in the interpreted *Se* and y-ions are in red.

### Alignment between experimental and theoretical spectra

Briefly, SpecGlobX like SpecGlob [19] relies on a dynamic programming technique to find the best alignment between the series of b-ion peaks in a theoretical spectrum *St* representing the amino acids of a peptide and pairs of peaks of the complemented spectrum *Se_c_*. SpecGlobX looks for an alignment that maximizes a user-defined score and possibly splits the mass difference ΔM between *St* and *Se* into several mass shifts.

We now detail the adaptations made on SpecGlob to implement SpecGlobX on experimental spectra and interpret them successfully, knowing that the filling of the dynamic programming matrix is described in detail in [19]. In SpecGlobX, the scoring system globally takes (positively) into account the number of aligned amino acids and (negatively) the number of inserted mass shifts and the number of amino acids that are not aligned. Each time SpecGlobX introduces a mass shift in an alignment (an elementary decision we call a realignment), this increases the risk to align a b-ion peak of *St* on an inappropriate ion peak of *Se_c_*. To mitigate this risk of misalignment, the score penalty is defined such that a realignment is preferred over non-aligned amino acids only if this realignment is subsequently compensated by the alignment of at least two amino acids. Thus, in the SpecGlobX scoring system, a realignment is more penalized than the non-alignment of one amino acid.

Considering the above rule, SpecGlobX uses the following scoring system during the dynamic programming step: (1) when a pair of peaks is aligned without any mass shift, the score is increased by 10 if the fragmentation generated both the b- and y-ion (illustrated in transitions 2 and 9 in Figure 1d) and by 7 otherwise (i.e., if only the b-ion or the y-ion peak is present in *Se*, as illustrated in transitions 1, 5 and 6 in Figure 1d); (2) when a pair of peaks corresponding to an amino acid is aligned, but this alignment requires a mass shift to displace the pair of peaks from an amino acid on a pair of peaks of *Se_c_*, then the score is decreased by 8 or 6 depending on whether the fragmentation generated both b- and y-ions or only one of the two (illustrated in transitions 4 and 8 Figure 1d); (3) when the amino acid is non-aligned, then the score is decreased by 4 (illustrated in transitions 3 and 7 in Figure 1d). The whole filling of the dynamic programming matrix corresponding to the alignment of the *St* and *Se_c_* spectra corresponding to the peptide HINATESVR is illustrated in Figure 1d. Lastly, SpecGlobX generates the best alignment under the form of a string as was done in [19] by a traceback step (Figure 1e).

For interested readers who want to dig deeper into the algorithm, we also mention a small subtlety that must be taken into account while the dynamic programming matrix is filled in SpecGlobX, compared to its previous version SpecGlob. A key observation in experimental spectra is that when a peak between two amino acids is missing in *Se_c_*, not only those two amino acids cannot be aligned (because their masses cannot be found in the spectrum), but this non-alignment generates a subsequent realignment whose mass shift is equal to a null value. To deal with this issue, this particular realignment (only due to one or several missing peaks) must be scored as an alignment (positive contribution to the score) rather than as a realignment (negative contribution to the score).

### Optimization of the location of the suggested mass shifts

In this last processing step, SpecGlobX adjusts the locations of the suggested mass shifts (if any) to maximize the number of shared peaks between *St* and *Se*. One of the most important goal of this step is to highlight the presence of a non-aligned mass if any.

The “completion process” compensates for a missing b-ion in *Se_c_* when the y-ion is displayed in *Se*, as long as the mass of the complementary peak can be directly inferred from the mass of the peptide. Regrettably, this relation is not always applicable, for instance when the precursor ion loses a neutral chemical group (such as water) in an early stage of the fragmentation process. Similar situations happen when *Se* merges the fragmentation of two peptides (possibly of identical sequences) linked by a chemical bond (such as a disulfide bridge) leading to homo or heterodimers. In these cases, all the complementary peaks are shifted from their expected value by a constant mass, which we denote as *Mloss*. Since the mass of the fragmented peptide observed in *Se* does not fit the measured mass of the experimental peptide, *Mloss* can be considered a non-aligned mass. To exemplify the generation of *Se_c_* in this instance, in Figure 2a, we display the spectrum we obtain after the completion process when a neutral loss of 301.99 Da (i.e. the mass of a frequent modification referenced in Unimod, although still annotated as ‘unknown’) has been added to the spectrum *Se* shown Figure 1a. In this example, Δ*M* is the sum of two mass shifts observed in *Se* (0.98 Da and 27.99 Da, *i.e*., the same mass shifts as in Figure 1) and one *Mloss* (301.99 Da). The dynamic programming process can then align some of the complementary peaks that do not correspond to any b-ion peaks in the original spectrum *Se*. The number of shared peaks between *Se_c_* and *St* is increased by this alignment, but the number of shared peaks between *Se* and *St* is not. This situation is illustrated by the simulated spectrum displayed in Figure 2. Since the b1-ion of peptide HINATESVR is not present in the spectrum, the mass of the first amino acid H is only given by the presence of the y8-ion. Then, the first amino acid H can be only aligned on the complementary peak shifted by 301.99 Da, a shift only due to *Mloss* in *Se_c_*. If we compute directly the number of shared peaks between *Se* and the suggested alignment [301.99]HI[N][0.98]ATE[S][27.99]VR, none of the y-ion peaks would align (due to *Mloss*). To circumvent this issue, the post-processing step evaluates whether each shift increases the number of peaks shared between *Se* and *St*. If not, the irrelevant shift is removed from the alignment and its value is accumulated to form the non-aligned mass indicated after the symbol ‘_’ in the result. We illustrate this transformation in Figure 2c.

**Figure 2.**
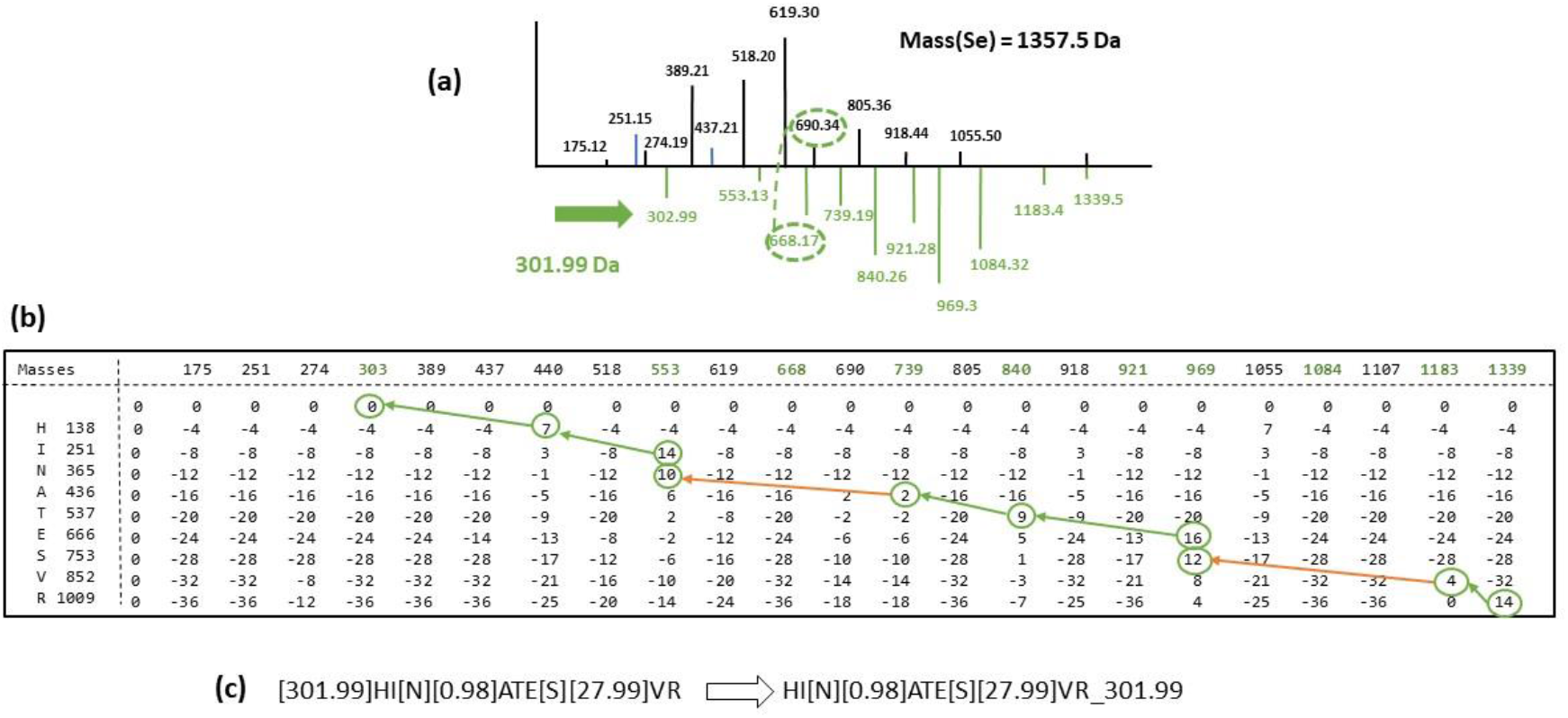
Alignment of peptide HINATESVR on a simulated spectrum carrying two modifications and a neutral loss. (a) The simulated spectrum *Se* is the same as the one presented in Figure 1a, except that the mass of the peptide has been increased by a neutral loss of 301.99 Da. As a result, complementary peaks in *Se_c_* –represented below the x-axis – are shifted by 301.99 Da compared to the complementary peaks displayed in Figure 1c; For example, the peak 690.34 is connected to its complementary peak 668.16 (366.17 + 301.99) by a dotted line (b) The score matrix computed by SpecGlobX to align the peptide HINATESVR with *Se_c_* by dynamic programming; Masses have been rounded for simplification of the graphical representation; (c) After the traceback step, *Se_c_* is interpreted as [301.99]HI[N][0.98]ATE[S][27.99]VR and after the post-processing step, *Se* is interpreted as HI[N][0.98]ATE[S][27.99]VR_301.99, identifying peptide HINATESVR with two modifications (a deamidation on “N” and a formylation on “S”) and a non-aligned mass of 301.99 Da (neutral loss), which is the correct identification.

In addition to bringing out the non-aligned mass, the post-processing step includes other transformations to facilitate the interpretation of alignments. For instance, non-aligned amino acids associated with a negative shift are removed if this deletion does not degrade the number of shared peaks (the case for a semi-tryptic peptide).

As a result of SpecGlobX, both alignments (before and after post-processing) are returned to the user.

## RESULTS AND DISCUSSION

### Application features

We implemented SpecGlobX as a standalone Java application (executable with example files are available in Additional file 1). As input, SpecGlobX reads a file containing spectra in MGF or mzML format (parsed by the JMZReader library [26]) and a comma-delimited spreadsheet that describes the list of PSMs to be aligned. This list of PSMs may be the result of any OMS method or a list of PSMs generated by a user.

The user has immediate access to the most frequent parameters from the interface (Figure 3), while he/she can also edit an additional file containing all the parameters and scores used in the implementation. A click on the ‘Launch Alignments’ button starts SpecGlobX as a mono or multi-thread execution and produces a comma-delimited CSV file as output. For each PSM, two alignments are returned: the ‘preAlignedPeptide’ column gives the alignment obtained just after the traceback step, while the ‘alignedPeptide’ column displays the alignment once the post-processing step is done. SpecGlobX returns all alignments as strings: amino acids not explained in the alignment and shifts are in square brackets. Therefore, the user visualizes which amino acids have corroborating peaks in the aligned spectrum. Moreover, the value of the non-aligned mass (if any) is written after the symbol ‘_’ at the end of the string. SpecGlobX also returns the number of peaks shared between *Se* and *St* before and after the alignments, and the percentage of the overall signal intensity explained by the alignment.

**Figure 3.**
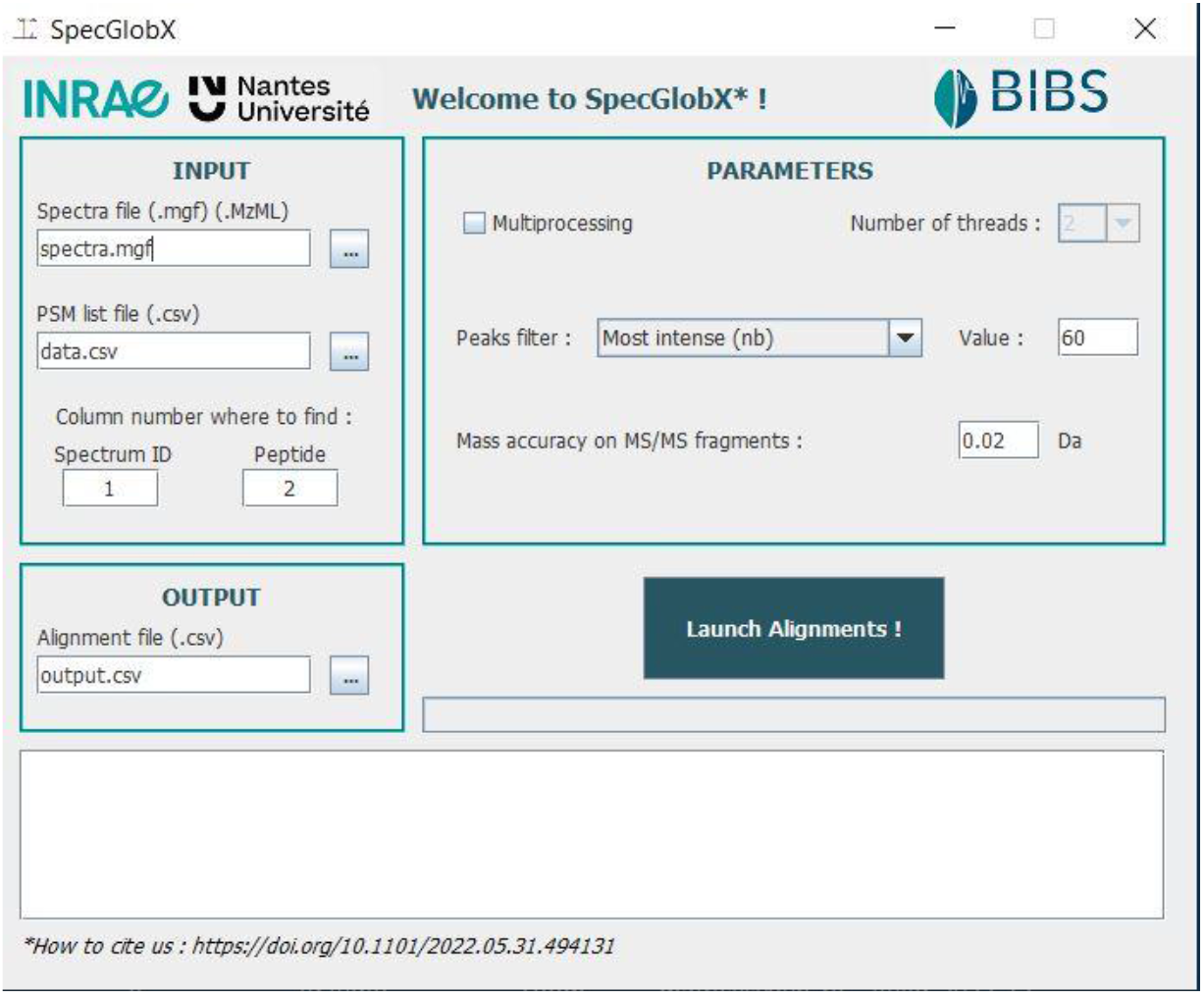
SpecGlobX graphic interface.

SpecGlobX is fast and can align one million PSMs in about 1.5 minutes on a desktop with 8 allocated threads and in about 10 minutes in a mono-thread configuration (around 100 times faster than InsPecT).

### Evaluation of SpecGlobX with a simulated spectra dataset

Firstly, we challenged SpecGlobX on a simulated spectra dataset *Dsim* generated from 50,000 tryptic unique peptides randomly selected from the human proteome - downloaded from Ensembl99, release GrCh38 [27] - and whose lengths span from 12 to 25 amino acids. We transformed those peptides into doubly charged simulated spectra with the following modifications: we introduced a deamidation on each asparagine (N+0.984016 Da) and a sodium adduct on each aspartic acid (D+21.981943 Da); next, we randomly removed 20% of the peaks to simulate missing peaks. Two-thirds of the removed peaks were b-ion peaks and one-third were y-ion peaks, as we observed that missing peaks are mainly b-ion peaks in experimental fragmentation spectra acquired in an HCD cell (a common instrument configuration of high-resolution mass spectrometers used in proteomics). Then, we simulated noise by adding up to 60 peaks of randomly selected masses. Lastly, we simulated a neutral loss of 17.03 Da on the peptide mass of each spectrum. We provided the complete list of ‘correct PSMs’ as input to SpecGlobX i.e., each simulated spectrum was associated with the peptide it derived from. Depending on the peptide’s composition, simulated spectra in *Dsim* contain a range of one to five modifications (among which at least one neutral loss of 17.03 Da). Given that every spectrum in *Dsim* is simulated, we can unambiguously determine whether a modification suggested by SpecGlobX is correct. Our assessment shows that SpecGlobX performs well: it detected the neutral loss on a large proportion of spectra (nearly 70%), and it correctly identified 53% of all the expected modifications (Table 1, first row).

**Table 1.** Performance of SpecGlobX measured on the simulated dataset *Dsim*. At first, we provided the full list of ‘correct PSMs’ as input to SpecGlobX: each simulated spectrum is associated with the peptide sequence it derives from (row 1). Then, we executed SpecGlobX on the PSMs returned by three different OMS methods (rows 2 to 4). Several percentages of correct identifications are summarized with and without (W/o) SpecGlobX. As a first criterion (column 2), a PSM is counted as correct if the spectrum is associated with the peptide sequence it derives from; the second criterion measures the percentage of PSMs that exhibits the neutral loss among the 50,000 PSMs (columns 3 and 4); the third criterion counts the PSMs in which all detected modifications are correctly identified and placed related to all PSMs (columns 5 et 6); the last criterion refers to the percentage of modifications that are correctly identified relative to the number of modifications (about 103,000) incorporated in the simulated spectra (columns 7 and 8).

| Origin of the PSMs | #Correct PSMs | % PSM identified with neutral loss |  | %PSM with all modifications correct |  | % correct modifications compared to expected |  |
| --- | --- | --- | --- | --- | --- | --- | --- |
|  |  | W/o SpecGlobX | With SpecGlobX | W/o SpecGlobX | With SpecGlobX | W/o SpecGlobX | With SpecGlobX |
| 'Theory' | 50 000 | NA | **68** | NA | **50** | NA | **53** |
| SpecOMS | 38 188 | 27 | **62** | 27 | **48** | 13 | **42** |
| MODPlus | 29 108 | 0 | **51** | 0 | **39** | 18 | **35** |
| MSFragger | 36 415 | 27 | **59** | 27 | **44** | 13 | **39** |

### Complementarity between SpecGlobX and existing OMS methods

Secondly, we evaluated whether SpecGlobX improves the results provided by three existing OMS software: MODPlus [14], MSFragger [9], and SpecOMS [23]. To this end, we first challenged each of these tools with *Dsim* spectra (see Supplementary Data, Additional file 2 for detailed parameters) and we examined each software’s ability to interpret the mass modifications in the correctly identified PSMs. On one hand, MODPlus explains ΔM observed between *Se* and *St* by combining modifications stored in a predefined list (consisting of all or only a selection of modifications referenced in Unimod, or a list provided by the user). We have previously shown that this strategy is particularly efficient if the objective is to identify the maximum number of peptides carrying common or known in-advance modifications. However, as soon as a peptide carries a modification not reported in the list, MODPlus fails in recovering a correct interpretation. Since MODPlus does not consider mass losses, it was unable to provide a good interpretation of any of the correctly identified PSMs, even though it interpreted and located 18% of the modifications (Table 1). On the other hand, MSFragger and SpecOMS try to explain each modification by a single shift. Therefore, both of them yielded good interpretations for only 27% of the correctly identified PSMs, which corresponded to peptides carrying a single modification (Table 1). Then, we applied SpecGlobX on the PSMs correctly identified by SpecOMS, MODPlus and MSFragger. Regardless of the search engine used to produce the PSMs, a large part of the neutral losses misinterpreted by the three search engines can be recovered by SpecGlobX. Similarly, the percentage of correct modifications relative to the number of expected modifications (based on the number of N and D in *Dsim*) is multiplied by two or three depending on the OMS software used. The limitation of SpecGlobX, however, is that even when the PSM is correct, only half of the modifications (53%) are perfectly identified and localized. This is not a surprise since some modifications cannot be corroborated without any *a priori* due to missing peaks. In addition, if two modifications are present on two adjacent amino acids, SpecGlobX is not able to distinguish them. Consequently, SpecGlobX can currently be viewed as a decision-support tool, particularly useful to highlight unexpected and complex modifications on peptides. A manual curation by an expert is still needed to remove ambiguities in the proposed alignments.

### Alignments highlighted on the HEK293 dataset

As a final evaluation, we ran SpecGlobX on a spectra dataset generated from HEK293 cells [24] downloaded from PRIDE (PXD001468). This spectra dataset has been used several times to evaluate open modification search methods. We converted the raw spectra file into the MGF format with msConvert release 3.0 [28].

To emphasize the value of SpecGlobX on experimental spectra, we focused on the interpretation of a subset of 77 spectra identifying peptide *pep*=DATNVGDEGGFAPNIIENK according to SpecOMS (results are in Additional file 3). This subset of spectra seems to be a good representation of a set of spectra identifying the same peptide in the dataset. Indeed, as shown in Figure 4, among the 77 spectra identifying *pep*, only 15 exhibit ΔM values around 0 or 1 Da (corresponding to an exact match to the expected mass or an incorrect picking of the good isotope, respectively), while the others are distributed over 44 different values of ΔM between −500 and 2065 Da. Surprisingly, when examining the alignments suggested by SpecGlobX, we note a large number of non-aligned masses (36 out of the 77 alignments returned by SpecGlobX contain a non-aligned mass), an observation we can extend to the alignments of all PSMs obtained on the complete dataset. Situations combining non-aligned masses including neutral losses and one or several modifications on the same peptide -as previously evaluated with *Dsim*-seems to be a reality in this dataset. The exploration of the spectra relations as done in [29] helps to decipher whether these neutral loss identifications are correct.

**Figure 4.**
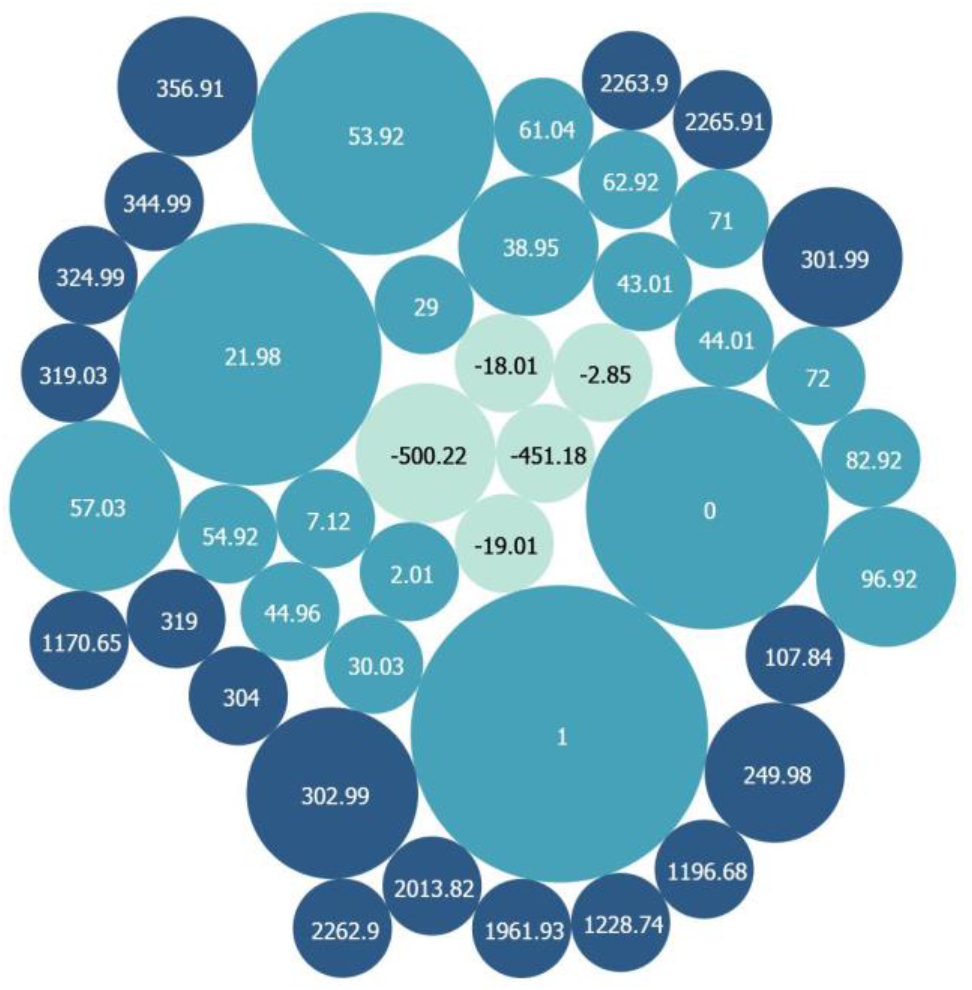
Representation of the variety of mass deltas carried by a set of spectra identifying peptide DATNVGDEGGFAPNIIENK (HEK293 dataset). The bubble areas, colored with a blue gradient according to the mass delta, are proportional to the number of spectra displaying each value of mass modification.

All the number of shared peaks between pairs of spectra are easily and rapidly completed by running SpecOMS (with the search_mode parameter set to “spectra”). Based on these data, we transformed the relations between the 77 spectra representing *pep* into a spectral network using GePhi [30], in which each node corresponds to a spectrum and two vertices are connected by a link when the two spectra share at least 26 out of the 50 most intense peaks. The resulting graph displayed in Figure 5 includes 74 connected nodes and 998 links. Links between pairs of spectra are weighted by their number of shared peaks. The force-directed layout algorithm Force Atlas [31] generated the detailed arrangement of nodes on the drawing. As a consequence, groups of nodes that are densely connected (i.e., that share the same set of peaks) are immediately highlighted as clusters of nodes.

**Figure 5.**
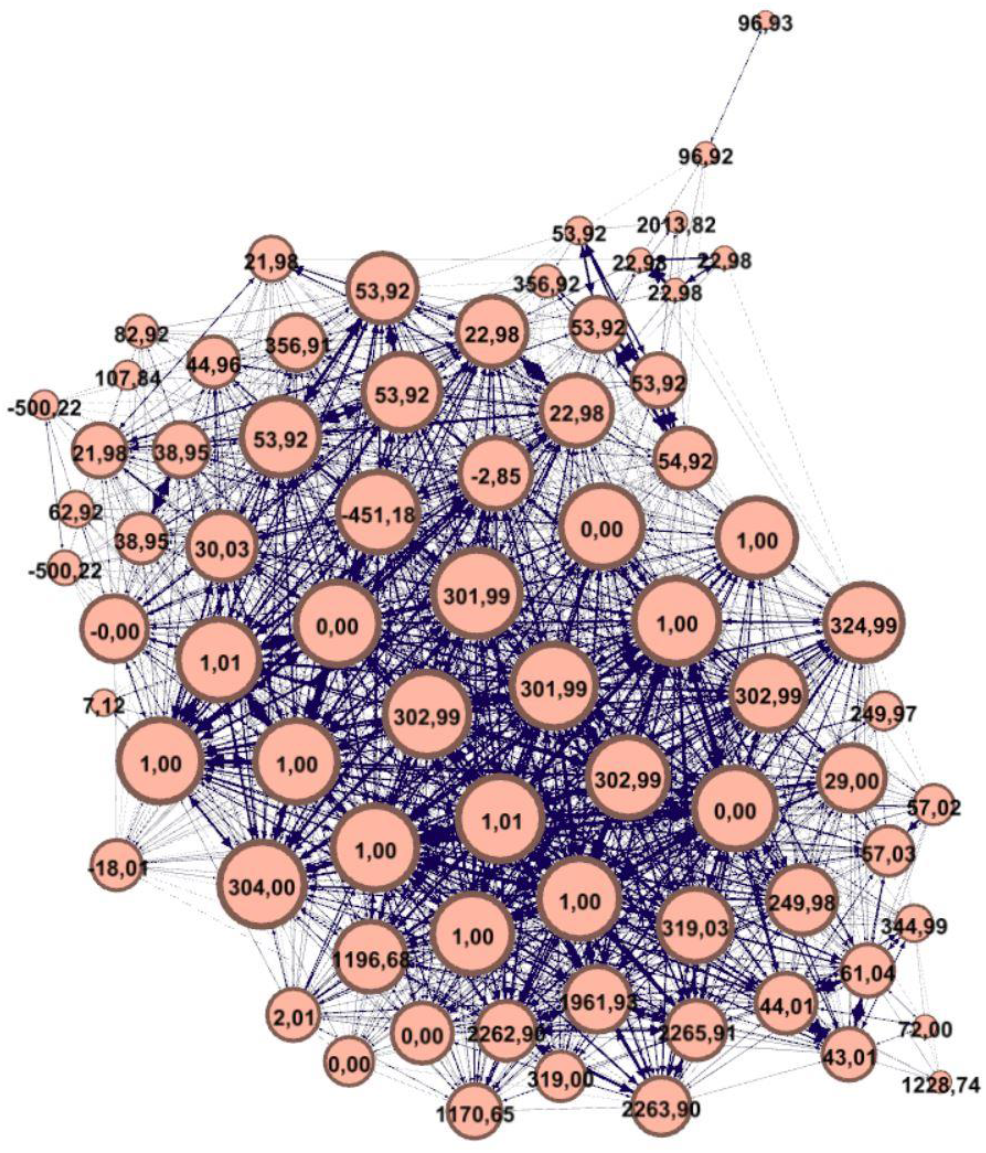
The closeness of spectra identifying peptide DATNVGDEGGFAPNIIENK (HEK293 dataset). In this picture, two vertices representing spectra are linked if the two spectra share at least 26 out of the 50 most intense peaks. The size of each node is related to its degree, the label on each node displays the ΔM value. Each link between a pair of spectra is weighted by the number of shared peaks. The arrangement of nodes is done by the directed Force Atlas algorithm so that the topology of the graph represents the closeness of spectra in terms of number of shared peaks.

We remind that two spectra representing *pep* with different neutral losses should share most of their peaks –and should share most of their peaks with the spectra identifying *pep* without modification (ΔM=0 or ΔM=1 corresponding to an incorrect use of isotopic peaks). On the contrary, the number of peaks shared between two spectra representing *pep* drops by half for each modification differentiating the two spectra. The presence of many non-aligned masses is confirmed by the topology of the graph with a large central cluster. For instance, ΔM= {301.99; 302.99; 303.99} Da most probably represent labile (and correlated) modifications. Consistent with that, all the spectra carrying these modifications and identifying *pep* with a strong confidence share most of their peaks with most of the other spectra, particularly with spectra associated with PSMs with ΔM = 0 or 1.

Next, we interpreted the 77 spectra identifying *pep* with MODPlus and MSFragger to evaluate if the behavior on experimental spectra is similar to what we observed on simulated spectra. The answer is clearly yes (the interpretation provided by each software is given in Additional file 3). Even though MODPlus provides the same PSMs as SpecOMS with only two exceptions, the interpretations in terms of modifications are the same for only 19 of these 75 PSMs. On the contrary, MSFragger differs more frequently on the PSM (5 spectra are interpreted as different peptides), but the interpretations of the modifications carried by 40 among 72 PSMs are the same.

As already highlighted by simulated spectra, well-known modifications are well-located by MODPlus (D+21.98 Da, N-Ter+43.01 Da, D+53.92 Da). However, one surprise is some of the nodes associated with the latter ΔM (Figure 5, ΔM = +53.92 Da for instance) are densely connected to nodes corresponding to mass losses, whereas some other nodes of the same ΔM are close to each other but disconnected from the cluster of nodes. This suggests that some spectra contain a clear signature of the modification with a shift of peaks carrying the modification, while other spectra have partly or totally lost this signature. These findings imply that non-aligned peaks are more frequent in spectra than previously thought, leading to a lower number of shared peaks if those peaks are not taken into account by search engines. Our small experimental dataset also illustrates that the counterpart of *a priori* knowledge and ΔM restrictions sometimes have bad effects on the interpretations of spectra. On one hand, spectrum #54025 interpreted without any doubt by MODPlus as the O-linked glycosylated peptide YGKDATNVGDEGGFAPNILENK (probability = 1, glycan 1914.69 Da) is more reliably interpreted as a homodimer of *pep* (an artifact that can be frequently observed [32]) added to the frequent neutral loss 301.98 Da with SpecGlobX and MSFragger. On the other hand, MSFragger suggests another peptide than *pep* for spectrum #53811, leading to an incorrect PSM. Indeed, some readers could be intrigued by the presence of spectrum #53811 associated with ΔM=-451.18 Da at the center of the cluster of nodes. The spectrum position in the graph suggests that #53811 shares most of its peaks with the entire peptide *pep* (nodes with ΔM = 0 or 1), which excludes the possibility that #53811 arises from a semi-tryptic form of ***pep*** (the MODPlus suggestion, but in this case, the number of shared peaks with other spectra representing *pep* would have been much lower). At the same time, the ΔM value is negative, suggesting a loss of amino acids. Even though none of the software has found this most probable interpretation, the usage of SpecGlobX as a decision support tool combined with an expert analysis of the spectrum graph has rather easily led us to the following explanation: the charge state evaluation of the spectrum is wrong (3+ rather than 2+) and the spectrum carries the labile modification of 302.99 Da (present on a large proportion of the spectra in this sample).

## Conclusions

SpecGlobX is an easy-to-use software developed to quickly align PSMs. SpecGlobX can be seen as an additional tool to existing OMS software, providing a good decision support tool to highlight complex modifications carried by peptides. We have shown on a large set of simulated spectra (modeled such as to simulate imperfect experimental spectra) that SpecGlobX improves the identification of complex and unanticipated modifications after reprocessing the PSMs lists provided by OMS software. Next, we demonstrated the usefulness of SpecGlobX on a subset of a well-known experimental spectra dataset downloaded from PRIDE, highlighting the presence of a large proportion of non-aligned masses due to neutral losses, charge estimation error, and presence of dimers.

SpecGlobX is freely available to all users, including the source code.

## Availability and requirements

The specGlobX project will be deposited (source + executable jar file) on GitHub after acceptance of the manuscript.

Requires java 1.8 or higher

## Availability of data and materials

This published article and its supplementary files give all information required to reproduce the simulated dataset *DSim*.

The HEK293 spectra dataset supporting the conclusions of this article was downloaded from the PRIDE repository, PXD001468, file b1922_293T_proteinID_02A_QE3_122212.raw

https://www.ebi.ac.uk/pride/archive/projects/PXD001468

## Acknowledgements

We express our sincere gratitude to Colette Larré, Michel Zivy, and Chantal Brossard for their help when we set up the deepProt project and for the enriching discussions we have shared.

## Supplementary information

### Additional file 1: SpecGlobX.zip

SpecGlobX.zip contains the SpecGlobX executable jar file, a user document (README.md), a default parameter file (config.properties), and a demonstration set of PSMs obtained by SpecOMS on the 77 spectra discussed in the article.

### Additional file 2: Parameters_ForEvaluation.pdf

Extended description of parameters used to process the spectra dataset *Dsim* by different software.

### Additional file 3: spectraInterpretation.xls

Comparative interpretation of a subset of spectra by three software

